# Quantitative mapping of pseudouridines in bacteria RNA

**DOI:** 10.1101/2024.11.26.625507

**Authors:** Shikha Sharma, Brendan Woodworth, Bin Yang, Ning Duan, Mannuku Pheko, Niki Moutsopoulos, Akintunde Emiola

**Affiliations:** Microbial Therapeutics Unit, National Institute of Dental and Craniofacial Research, National Institutes of Health, Bethesda, Maryland, United States of America; Human Barrier Immunity Section, Laboratory of Host Immunity and Microbiome, National Institute of Allergy and Infectious Diseases, National Institutes of Health, Bethesda, Maryland, United States of America

## Abstract

RNA pseudouridylation is one of the most prevalent post-transcriptional modifications, occurring universally across all organisms. Although pseudouridines have been extensively studied in bacterial tRNAs and rRNAs, their presence and role in bacterial mRNA remain poorly characterized. Here, we used a bisulfite-based sequencing approach to provide a comprehensive and quantitative measurement of bacteria pseudouridines. As a proof of concept in *E.* coli, we identified 1,954 high-confidence sites in 1,331 transcripts, covering almost 30% of the transcriptome. Furthermore, pseudouridine mapping enabled the detection of differentially expressed genes associated with stress response that were unidentified using conventional RNA-seq approach. We also demonstrate that in addition to pseudouridine profiling, our approach can facilitate the discovery of previously unidentified transcripts. As an example, we identified a small RNA transcribed from the antisense strand of tRNA-Tyr which represses expression of distal genes. Finally, we mapped pseudouridines in oral microbiome samples of human subjects, demonstrating the broad applicability of our approach in complex microbiomes. Altogether, our work highlights the advantages of mapping bacterial pseudouridines and provides a tool to study posttranscription regulation in microbial communities.

## Introduction

Pseudouridines (Ψs) are one of the most abundant nucleotide modifications present in all domains of life^1,2^. They are found in rRNA, tRNA, and other non-coding RNA where they enhance base-pairing, RNA stability, and influence translation fidelity^3,4^. Recent transcriptome-wide studies in humans have further identified Ψs in eukaryotic mRNAs where pseudouridylation is altered in response to stress which suggests a regulatory role in eukaryotes^5^.

In many bacteria species such as *E. coli*, modification is carried out by eleven different pseudouridine synthase (PUS) enzymes ─ five of which specifically modify rRNA targets (RluB, RluC, RluD, RluE, and RsuA)^6,7^. On the other hand, TruA, TruB, TruC, and TruD explicitly modify tRNAs, while RluA and RluF pseudouridylates both rRNA and tRNA^6^. Although, lack of Ψ in eukaryotic rRNA severely impacts ribosomal function, mutant *E. coli* strain devoid of any Ψ in the ribosomal subunit did not show any major effect on growth, decoding and ribosome biogenesis^7^. Among the tRNA PUS enzymes, TruA appears to be the most important. Mutations in TruA, which is responsible for pseudouridylation in anticodon loop, affects the translation machinery^8^. In particular, tRNA lacking Ψs in this loop, are unable to proceed at aminoacyl-tRNA transfer step due to lack of stability in the mRNA-tRNA complex^8^.

Most *in vivo* studies of Ψ in bacteria have been focused on rRNA and tRNA. This is partly due to the unavailability of experimental toolset to investigate the distribution and function of Ψ in mRNA. Previous *in vitro* studies have demonstrated that replacement of uridine with Ψ in bacteria mRNA can impede amino acid addition and increase the occurrence of amino acid substitutions^9^. Similarly, Ψ at stop codons (UAA, UAG, or UGA) can enable ribosomal readthrough of the modified stop codon^10^. Based on these observations, Ψs in mRNAs are believed to influence protein translation in bacteria, but yet to be investigated *in vivo*.

There has been one attempt to identify Ψ sites in bacteria mRNA at base-level resolution^11^. The authors relied on Pseudo-seq, a transcription-wide approach for Ψ profiling^12^. However, Pseudo-seq is known to suffer from low sensitivity and typically identifies very few Ψ sites. In fact, newer methods that uses bisulfite (BS) to label Ψ (e.g. BID-seq^13,14^ and PRAISE^15^) have recently been shown to identify ∼23 times more Ψ sites than Pseudo-seq^14^. Therefore, new approaches are required to evaluate the widespread distribution of Ψ in bacteria mRNAs.

Furthermore, studying mRNA pseudouridylation in microbiomes may help elucidate the role of many genes under a given condition. Typically, analysis of gene expression of the microbiota is performed using RNA-seq (i.e. metatranscriptomics). In this case, mRNA level in sequence data is used as an indirect proxy for protein abundance. Given that Ψ a common mRNA modification and its potential effect in altering translation^13^, conventional RNA-seq analysis may not accurately recapitulate the protein translation landscape of the microbial community. Consequently, new methods capable of incorporating this widespread posttranscription modification will provide a more accurate representation of protein dynamics. To date, there is no study that has examined mRNA pseudouridylation or posttranscriptional regulation in microbiomes.

In this work, we adapted a BS-based approach to identify Ψ sites in *E. coli* RNA at base resolution. We identified Ψ sites in one-third of mRNAs and established that Ψ profiling can identify transcripts associated with stress conditions which are undetected using conventional RNA-seq. We also establish that novel transcription units can be identified using Ψ profiling. Lastly, we demonstrate the applicability of our method in complex microbiome samples. Altogether, our work provides a tool to study posttranscription regulation in isolate bacteria and complex microbial communities.

## Results

### A BS-based approach to identify Ψ positions in bacteria RNA

BS-based approaches were recently developed to identify the positions of Ψs in human mRNA^13–15^. However, due to the instability of prokaryotic mRNAs and absence of poly-A tails, these methods are not readily applicable to bacteria. In BS methods, treatment of mRNA with BS generates BS-Ψ adduct, which subsequently induces deletion at Ψ sites during reverse transcription (RT). By mapping these single base deletions with untreated samples (which have no deletions), we can identify the exact location of Ψs (Fig. 1). We adapted the BS-based approach with modifications and applied it to *E. coli* samples.

**Figure 1:**
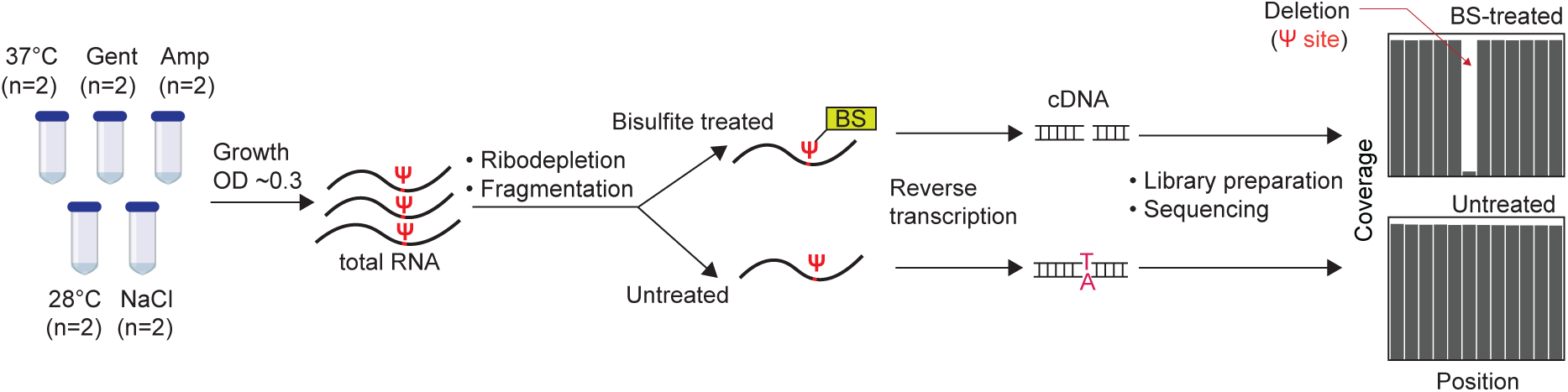
Pipeline for detection of Ψ sites in bacteria RNA. BS treatment of RNA induces deletion at Ψ locations during cDNA synthesis. After sequencing, deletion sites in BS-treated samples are compared with untreated samples to accurately identify Ψ positions. Gent = gentamicin; Amp = ampicillin.

Briefly, we began by extracting total RNA and enriching for mRNA through ribodepletion. Next, we performed fragmentation and split samples into two halves, with one half treated with BS (Fig. 1). After RT and sequencing, we mapped reads to the *E. coli* genome, realigned reads to increase sensitivity for reads that contain gaps^16^ and retrieved the coverage of each nucleotide. Based on thresholds from previous studies^13,15^, we considered a uridine position as pseudouridylated if (i) the coverage depth in both BS-treated and untreated samples is ≥ 20; (ii) deletion rate is ≥ 5% in BS-treated samples but less than 1% in untreated samples; and (iii) deletion count is above 5 in BS-treated samples. Therefore, the deletion fraction is proportional to Ψ levels. For instance, a deletion fraction of 0.5 suggests 50% of transcripts harbor Ψ in a site.

### Validation of BS-based profiling of bacterial Ψs

To validate our approach, we used a wild-type (WT) *E. coli* and a mutant strain with knockout deletions in all seven rRNA PUS enzymes^7^ (*rluA, -B, -C, -D, -E, -F,* and *rsuA*). This mutant is incapable of pseudouridylating 16S and 23S rRNA and subsequently referred to as Ψ^ΔrRNA^ strain. We cultured cells under optimum (37°C) and various stress conditions (28°C, ampicillin, gentamicin, and high NaCl) to maximize the detection of Ψ sites, especially in transcripts which are mostly expressed under sub-optimal conditions (Extended Data Fig. 1A). Using the pipeline described above, we initially sought to determine the distribution of all base deletions in our dataset. Because the GC content of *E. coli* is approximately 50% and assuming deletions occur randomly, the expected deletion counts for a given base, relative to other bases, is expected to be similar under non-treated conditions. In other words, the average deletion ratio for one base (e.g. U), relative to others should be ∼ 1. (i.e. mean(*U:A, U:G, U:C)*). Indeed, in untreated samples, we observed the expected deletion frequencies for each base (Fig. 2A). In contrast, U-sites, but not other bases, were deleted > 5 times above the expected frequency in BS-treated samples. This is consistent with the hypothesis that BS-treatment results in deletions at Ψ positions^13–15^.

**Figure 2:**
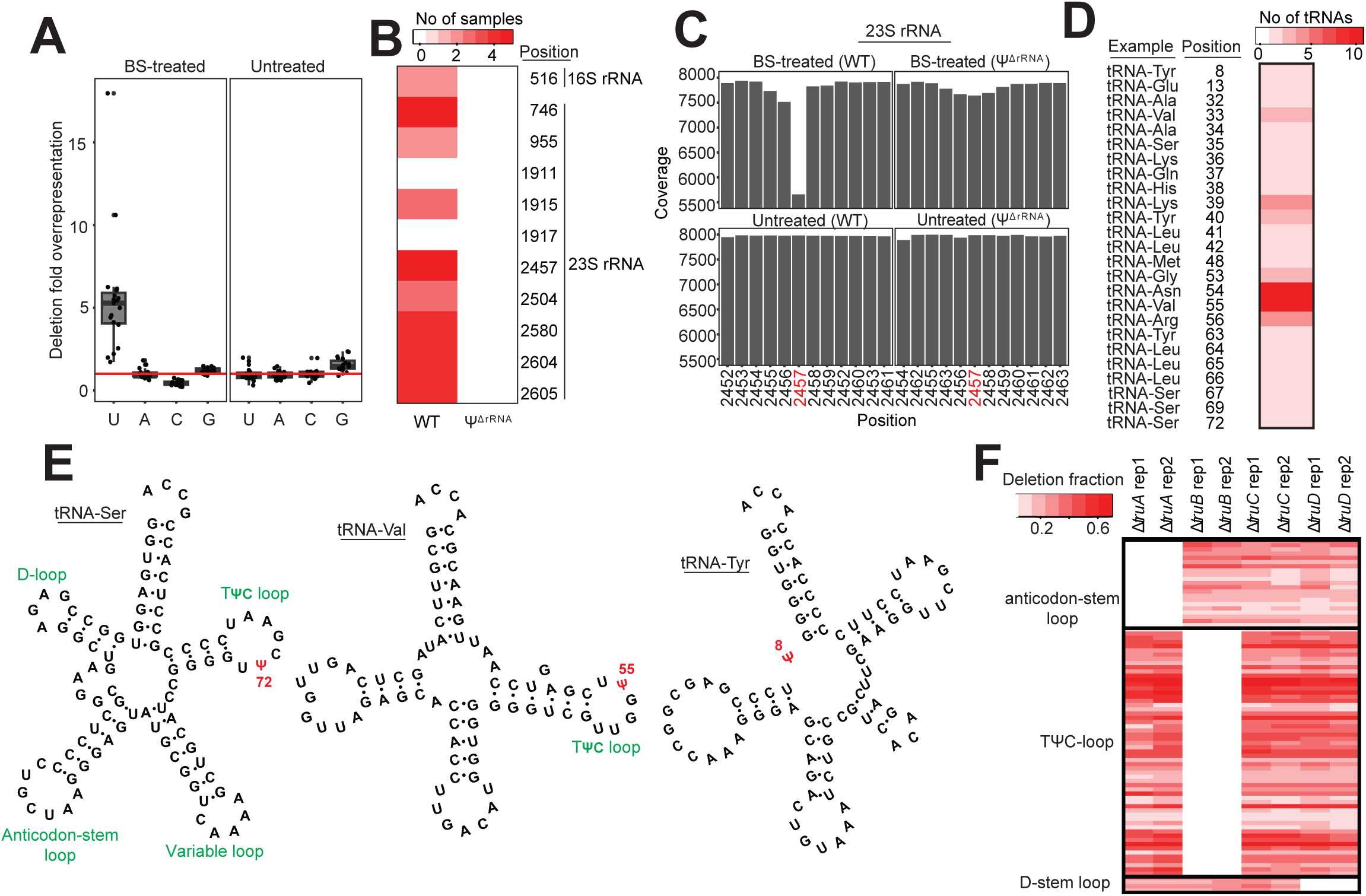
Bisulfite-based profiling accurately detects known Ψ sites in bacteria rRNA and tRNA. (A) Deletion frequency of each nucleotide under BS-treated and untreated conditions. Since the GC content of *E. coli* is approximately 50%, random deletions, relative to other bases should be similar. The horizontal line represents the expected deletion frequency if deletion occurs randomly. BS-treatment induces deletions specifically at uridine sites. (B) Detection of known Ψ positions in rRNA from WT and a mutant strain unable to pseudouridylate rRNA (Ψ^ΔrRNA^). The heatmap shows the number of samples where Ψ was detected. (C) Representative view of a Ψ site (position 2,457) in 23S rRNA from WT and Ψ^ΔrRNA^ strains. (D) Identification of Ψ sites in tRNAs. The heatmap shows the number of tRNAs with detected Ψ at a given site. Representative examples of tRNA harboring Ψ modification in a particular site is also provided. (E) Examples of tRNA structures (tRNA-Ser and tRNA-Val) harboring Ψ in exact locations in the TΨC loop despite having varying length. The structure on the right (tRNA-Tyr) shows a newly identified Ψ site in position 8. (F) Ψ proportion (deletion fraction) from biological replicates of tRNA PUS mutants. Each row represents a Ψ site in tRNAs.

For downstream analysis, we included an additional filter to minimize the detection of false positives. We required a putative Ψ site to be identified in at least two samples ─ whether from biological replicates or different stress conditions. This resulted in 1,954 high-confidence sites in 1,331 transcripts, covering almost 30% of the transcriptome. Importantly, the deletion rates were highly reproducible across biological replicates, indicating a quantitative ability of BS-based methods to detect Ψ in bacteria mRNAs (Extended Data Fig. 1B, Supplementary Table 1). To further validate our approach, we searched for known Ψ sites in rRNA. Although all samples were subjected to ribodepletion during sample prep (Fig. 1), there is usually a small percentage (∼ 0.5%) remaining which is sufficient for many analyses^17^. Our pipeline accurately identified the sole Ψ site in 16S rRNA, and 8 of 10 sites in 23S rRNA (Fig. 2B and C). In contrast, we did not detect any Ψ in rRNAs from Ψ^ΔrRNA^ samples (Fig. 2B and C). Therefore, our approach accurately identified known Ψ positions in rRNA with no false positives detected.

Next, we sought to identify established Ψ sites in tRNAs. Pseudouridines are known to be deposited in position 13 (D-stem loop), positions 32, 38, 39, and 40 in the anticodon-stem loop, and positions 55 and 65 in the TΨC-loop^6,18,19^. Similarly to the rRNA data above, we accurately detected all known tRNA sites (Fig. 2D), despite differences in length due to the variable arm region. For instance, with our pipeline, we found position 72 in the TΨC loop of tRNA-Ser is pseudouridylated, which is equivalent to position 55 in the TΨC loop of tRNA-Val (Fig. 2E). Furthermore, we uncovered a previously unreported Ψ site in position 8 of tRNA-Tyr (Fig. 2E). This is unlikely to be a false positive since the sequence motif (GUUC) is similar to that found in TΨC loops^13,15^ (Fig. 2E).

Lastly, we obtained four different PUS knockout strains that are incapable of pseudouridylating specific positions in tRNAs. We found Δ*truA* strains were unable to deposit Ψ in the anticodon-stem loop (Fig. 2F). Similarly, we did not detect Ψs in TΨC-loop and D-stem loop of tRNAs in Δ*truB* and Δ*truD* strains, respectively (Fig. 2F, Supplementary Table 2). These observations are consistent with established target locations of TruA, TruB, and TruD PUS enzymes^18,19^. However, although TruC modifies position 65 in the TΨC-loop, we still observed Ψ at this location in Δ*truC* mutant (Fig. 2F). One possible explanation could be low efficiency of TruC *in vivo* since it is known to modify only two tRNAs^20^. Alternatively, TruB may deposit Ψ in TruC sites since they recognize similar motif as described below. Altogether, our approach accurately identifies known Ψ positions in rRNA and tRNAs of *E. coli*.

### A comprehensive Ψ landscape in *E. coli* transcriptome

In a previous study to map the repertoire of Ψ in *E. coli* RNAs, Schaening-Burgos *et al* identified Ψ in only 42 mRNAs^11^ using Pseudo-seq. However, using the highly sensitive BS-based approach, we detected high-confidence Ψ sites in 1,217 mRNAs, which is 27 times above previous estimates (Fig. 3A, Supplementary Table 1). Most mRNAs only harbor a single Ψ site (Extended Data Fig. 2A). In addition, we found few mRNAs that are always pseudouridylated irrespective of growth conditions. For example, the multidrug efflux transporter, *mdtG,* contained Ψs in both normal and all tested stress conditions (Fig. 3B). In addition, almost 20% of detected pseudouridylated mRNAs are either involved in the biosynthesis of secondary metabolites or adaptation to diverse environments (Fig. 3C). This suggests a possible regulatory role of Ψ in response to stress, similarly to eukaryotes^5^.

**Figure 3:**
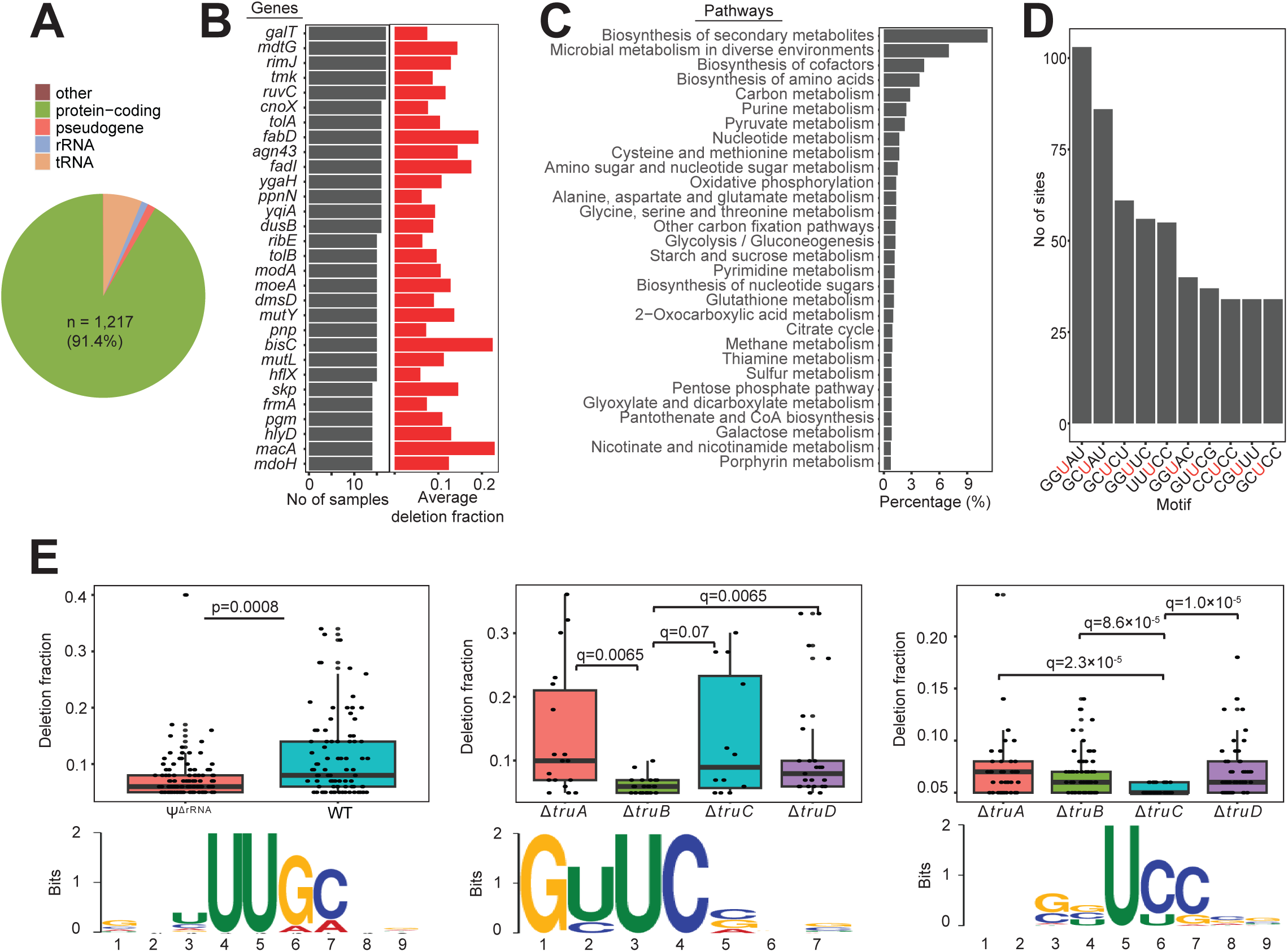
Quantitative landscape of Ψ in *E. coli* transcriptome. (A) Pie chart showing the distribution of Ψ across RNA types. (B) Ψ profiles of top 30 mRNAs. The barplot on the right shows the average deletion fraction across samples. For an mRNA with multiple Ψ positions, the sum of deletion fractions at all Ψ sites was calculated. (C) Pathway representation of mRNAs with Ψ. (D) Distribution of the top 10 sequence motifs for pseudouridylation. The red uridine residue represents the modification site. (E) Motifs with decreased deletion fractions in Ψ^ΔrRNA^ (left), Δ*truB* (middle), and Δ*truC* (right) strains, respectively. The image below shows the respective consensus recognition motifs. Significant differences between groups were computed with two-sided Wilcoxon rank sum test (left) or two-sided Wilcoxon rank sum tests with false discovery rate (FDR) adjusted *P* values (*q*).

Next, we analyzed the motif frequency and distribution of all Ψ sites. The top motifs contained mostly G or C bases upstream of the Ψ site (Fig. 3D, Supplementary Table 3). Interestingly, the most abundant motif (GGUAU), which we detected in over 100 sites, is also reported as being a frequent Ψ motif in the human transcriptome^15^. To further understand the sequence preference for each PUS enzyme, we compared the deletion fraction for every motif in WT and mutant strains. In other words, a motif with a significantly lower deletion rate in a PUS mutant is most likely a sequence preference. We established that target sites for rRNA PUS enzymes occur in a consensus sequence comprised of ‘UUGC’ (Fig. 3E) which agrees with previously reported RluA recognition motif^11^. Similarly, we observed ‘GUUC’ as the main recognition sequence for TruB, in strong agreement with the homologous TRUB1 targets in human and yeast^13,21^ (Fig. 3E). TruC, on the other hand, deposits Ψ mostly in ‘UCC’ sequences (Fig. 3E). However, we did not identify a consensus sequence for TruA or TruD. This may suggest a lack of sequence preference or pseudouridylation by other PUS. We also examined the secondary structure of all target sites and observed pseudouridylation occurs predominantly in unpaired uridine sites (Extended Data Fig. 2B). However, deletion rates were similar in both paired and unpaired sites (Extended Data Fig. 2C). Overall, Ψs are widespread across *E. coli* mRNAs and are mostly deposited in loop or hairpin structures by rRNA and tRNA PUS enzymes.

### Ψ stabilizes bacterial mRNA

Because Ψs are strongly linked to stability of non-coding RNA^22,23^, we examined their possible role in mRNA stability and gene expression. To begin, we retrieved transcripts containing Ψ that can be assigned unambiguously to a PUS enzyme. In this case, positions with lower deletion fraction in a PUS mutant, relative to WT, was considered a recognition site for a given PUS. Since multiple PUS can deposit Ψ at different locations in a single transcript, we assigned those candidates to multiple PUS. Next, we determined the abundance levels of candidate mRNAs by calculating the ‘transcript per million’ (TPM). We observed that mRNAs with lower deletion fraction were significantly less abundant (Fig. 4A). This was true for all PUS targets, with the sole exception of TruA. Taken together, our results underscore a functional role of pseudouridylation in stabilizing bacteria mRNA.

**Figure 4:**
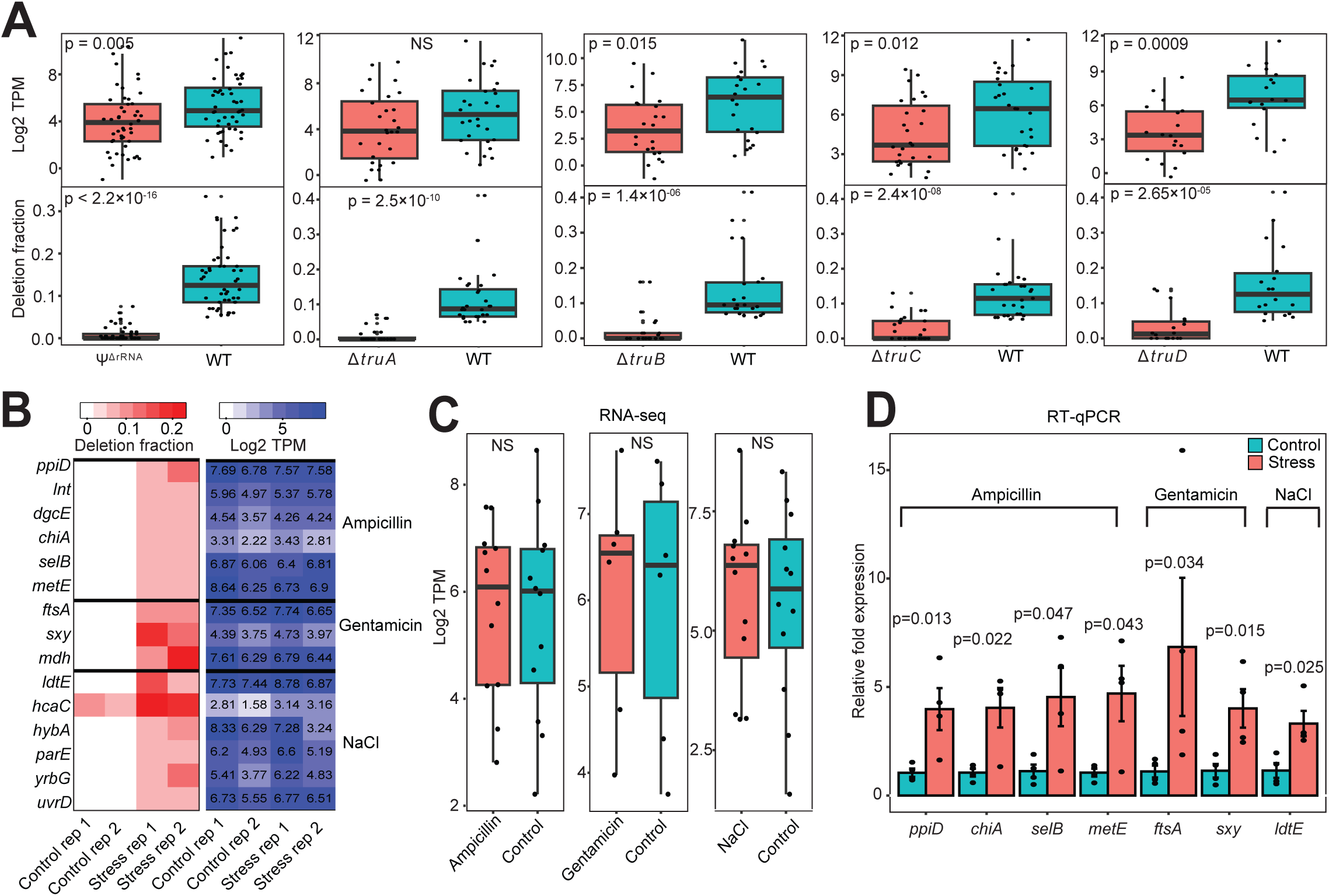
Ψ is associated with mRNA abundance. (A) Boxplot showing mRNA abundance (TPM) in WT and PUS mutants (top). Each datapoint represents the mean from 2 biological replicates. The lower boxplot shows the corresponding deletion fraction. Significant differences between groups were computed with two-sided Wilcoxon rank sum test. NS, not significant. (B) Heatmap showing mRNAs from WT cells with increased Ψ (deletion fraction) under ampicillin, gentamicin, or NaCl stress conditions from two biological replicates. The heatmap on the right shows the corresponding mRNA abundance (TPM) from RNA-seq. (C) Boxplot showing mRNA abundance (TPM) under control and stress conditions using data from ‘B’ above. Each data point represents the mean from two biological replicates. Statistical tests were performed using two-sided Wilcoxon rank sum test. NS, not significant. (D) Relative levels of mRNAs significantly enriched under stress conditions from four biological replicates and the error bars represent standard error of mean (SEM). Statistical significance was determined from normalized *Ct* values using unpaired, two-tailed *t*-test.

### Ψ profiling identifies differentially expressed genes not captured by conventional RNA-seq

Since pseudouridylation has a regulatory function in eukaryotes^5^, we sought to identify mRNAs with increased Ψ in response to ampicillin, gentamicin, and high salinity stress conditions. Among the identified candidates, 14 of 15 mRNAs had no detectable Ψ in control samples (37°C) (Fig. 4B). Interestingly, some of these candidates are known to be specifically associated with the treated stress condition. For example, *mdh* plays a role in resistance to gentamicin^24^, while *yrbG* is a Na^+^/Ca^2+^ cation antiporter that contributes to cellular adaptation in saline environments^25^. Because we already established that Ψ is linked with mRNA abundance, we hypothesized that the observed increment in pseudouridylation may be due to elevated mRNA under stress conditions. Surprisingly, we found no difference in expression levels in control and stress samples using TPM estimations (Fig. 4B and C). To further verify our hypothesis, we performed RT-qPCR to quantify transcript abundance. Indeed, we found seven candidates were significantly abundant under stress conditions (Fig. 4D). For candidates that failed to meet the significance threshold, we also observed a trend towards stability under these stress conditions (Extended Data Fig. 3). Altogether, Ψ profiling can facilitate the discovery of genes associated with bacterial adaptation to varying environmental conditions which may not be identified using other sequencing-based methods.

### Discovery of novel transcription units using Ψ profiling

Due to the complexity of bacterial transcriptomes, it is believed there are many transcription units (TUs) yet to be uncovered, even for well-studied model organisms such as *E. coli*^26^. In addition to Ψ mapping, we hypothesized our pipeline can identify novel TUs, especially antisense RNAs (asRNAs). To begin, we searched the opposite strand of all genes for presence of Ψ and only considered candidates detected in at least two samples. In total, we identified 258 high-confidence Ψ sites in the reverse strand of 246 genes which strongly suggests the expression of asRNAs (Supplementary Table 4). To validate our findings, we examined the proximity of identified Ψ sites to the transcription start sites (TSS) of published asRNAs^27^. Indeed, over 26% of Ψ positions were found within 200 bp, downstream of known TSS. This suggests the identified Ψs are likely positioned within the antisense transcript (Fig. 5A).

**Figure 5:**
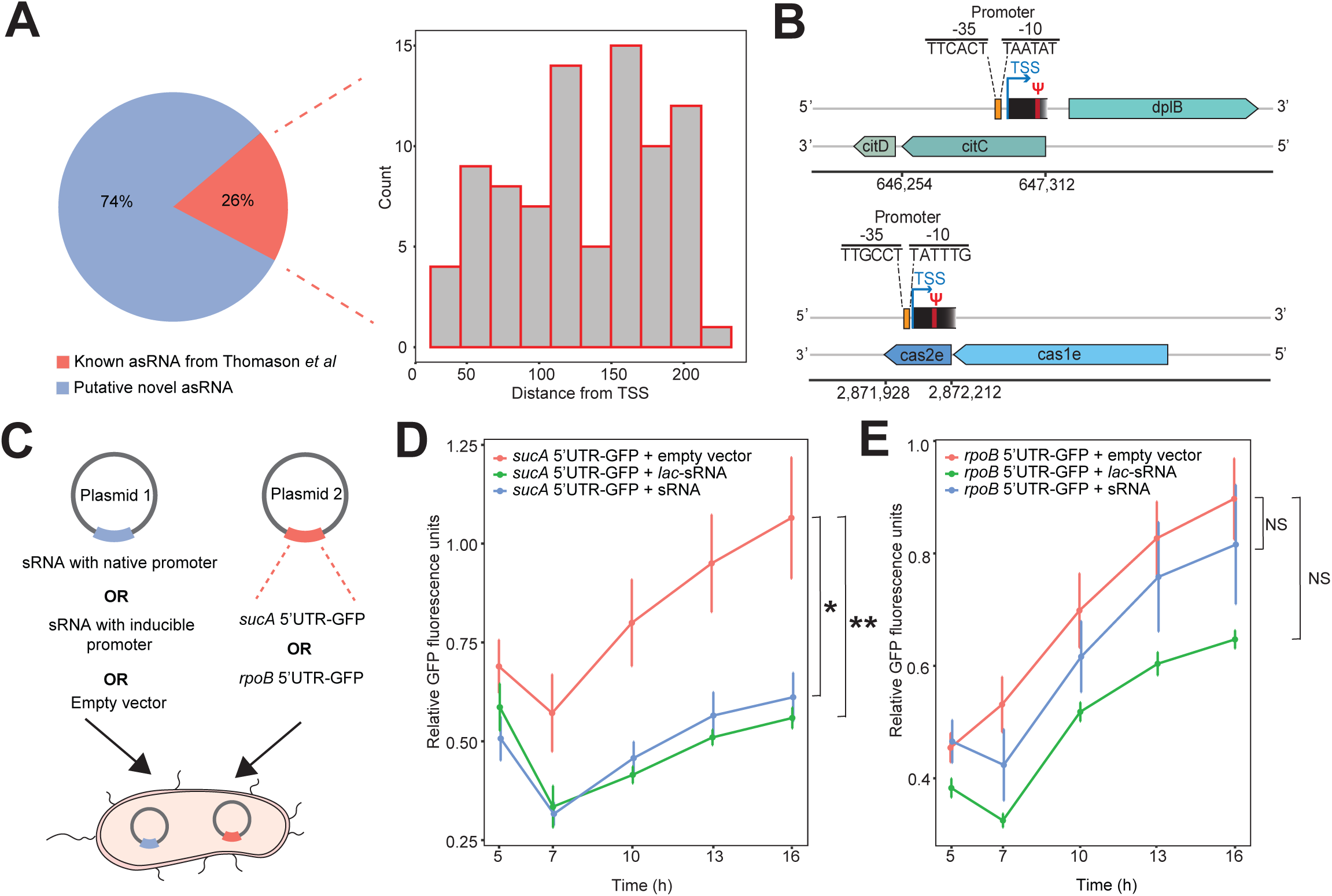
Ψ profiling can facilitate discovery of novel transcripts. (A) Pie chart showing the proportion of predicted asRNAs that matches previous results from Thomason *et al.* The majority of putative asRNAs are novel. The histogram shows the distance distribution between Ψ locations and transcription start sites reported in Thomason *et al*. (B) Examples of putative novel RNAs transcribed from the antisense strand of *citC* and *cas2e* genes. (C) Schematic representation of double-plasmid expression system to examine gene repression by sRNA. (D) Relative expression of GFP (fluorescence) over time in cells containing a *sucA*-5’UTR conjugated GFP plasmid. Raw fluorescence values were normalized to OD_600_ and timepoint 0. Experiments were conducted in M9 minimal media using five biological replicates and error bars represent standard error of mean (SEM). *P < 0.05; **P < 0.01, by one-way analysis of variance (ANOVA) with Tukey’s multiple comparison test at timepoint 16 h. *lac*-sRNA = sRNA with lac inducible promoter; sRNA = sRNA with native promoter. (E) Relative fluorescence in cells harboring a *rpoB*-5’UTR conjugated GFP plasmid. NS, not significant.

Though, majority (74%) of the putative asRNA identified in our data are putatively novel, they are unlikely to be false positives partly because prior studies relied on bacteria cultures grown in rich or minimal media^27^ whereas, we analyzed samples from multiple different stress conditions. To further confirm our results, we searched for promoter sequences upstream of Ψ locations via the software iProEP^28^. Using a very strict probability cut-off (> 0.9), 106 of 186 sites (57%) had an upstream promoter sequence within 200 bp (Fig. 5B). Consequently, our candidates likely represent bona fide asRNAs.

Another advantage of our approach is that it can help identify new small RNAs (sRNAs) that may play important regulatory roles in maintaining cellular homeostasis. In candidates where we identified an upstream TSS, we similarly searched for transcription termination sites (TTS) downstream of the Ψ site. In other words, we focused on ∼ 350 bp sequences harboring a promoter region, Ψ site, and a TTS. Overall, we identified 4 putative antisense sRNAs with predicted lengths ranging from 66 to 171 bp (Extended Data Fig. 4A, Supplementary Table 5). One candidate putatively transcribed from the antisense strand of tRNA-Tyr was of interest because, unlike others, we computationally predicted it may target multiple different mRNAs by binding to their 5’ untranslated region (UTR) to repress translation (Extended Data Fig. 4B, Supplementary Table 6). To test our hypothesis, we selected two targets ─ one predicted to be weakly inhibited (*rpoB*) and the other, strongly repressed (*sucA*). We used a double-plasmid expression system where one plasmid contained the sRNA sequence under the control of its native promoter or a *lac* inducible promoter. In the second plasmid, the 5’UTR of predicted targets was directly linked to the coding sequence of green fluorescent protein (GFP) (Fig. 5C, Extended Data Fig. 4C, Supplementary Table 6). After transformation and growth in M9 minimal media, we found a significant reduction in fluorescence levels in cells containing a *sucA*-5’UTR conjugated GFP plasmid (Fig. 5D). The findings were reproducible using sRNA expressed either under the control of its native promoter or induced (Fig. 5D). On the other hand, we did not observe repression in cells harboring a *rpoB*-5’UTR linked GFP plasmid (Fig. 5E). Thus, our candidate potentially inhibits the translation of *sucA in vivo.* Taken together, Ψ profiling can enhance the discovery of previously unidentified transcripts that can play important roles in regulating gene expression.

### Quantitative mapping of Ψ landscape in human microbiome samples

To examine the ability of our approach to quantify Ψ in complex microbial communities, we recruited two patients with periodontitis and retrieved oral plaque samples from the subgingival region (Fig. 6A). After BS treatment and sequencing, we mapped reads to approximately 2 million non-redundant genes retrieved from publicly available oral metagenomes^29^ (Fig. 6A). Unlike our approach for *E. coli* isolates where we required a Ψ site to have a deletion rate < 1% in untreated samples, we considered sites with higher deletions to accommodate indels arising from strain heterogeneity in complex microbial communities. However, we required the deletion rate in BS-treated samples to be greater than twofold in untreated samples. To further minimize false positives, we used a *P* value-based approach which takes into consideration total read depth, deletion rate, and relationship between these parameters in BS-treated and untreated samples^14^.

**Figure 6:**
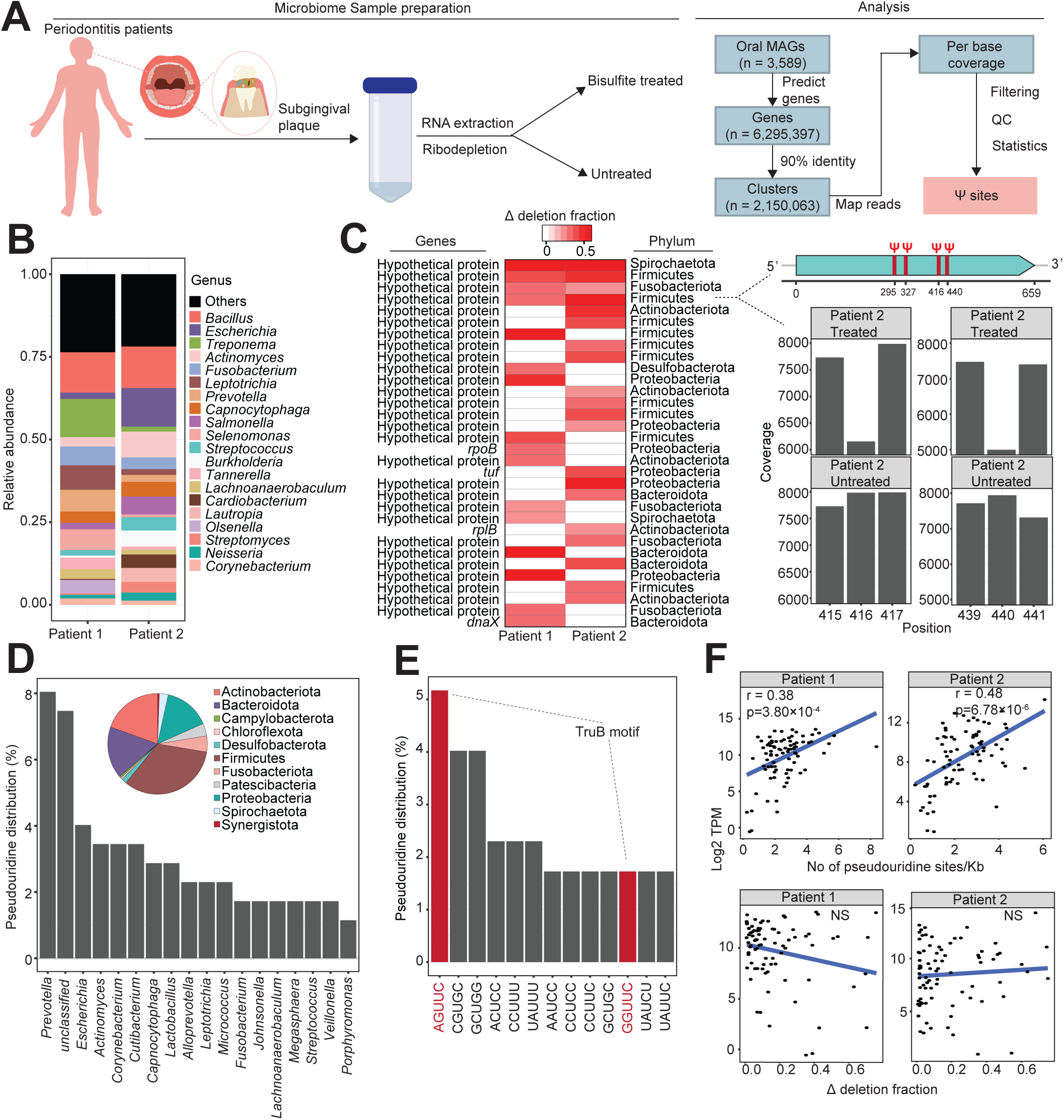
Quantitative mapping of Ψ in the oral microbiome. (A) Experimental and computational pipeline to process oral microbiome samples. (B) Microbial relative abundance from two periodontitis patients. (C) Top pseudouridylated mRNAs in oral samples. Ψ proportion is measured as the difference in deletion fraction between BS-treated and untreated sites (i.e. Δ deletion fraction). The image on the right shows an example of an uncharacterized Firmicutes mRNA with four different Ψ sites. The barplot is a representative coverage profile of Ψ sites (positions 416 and 440) in the mRNA above. (D) Proportion of pseudouridylated mRNAs across the top bacteria genera. The inset pie chart shows distribution across all phyla. (E) Distribution of top sequence motifs for pseudouridylation. The red bars represent the ‘GUUC’ recognition motif in TruB-dependent sites. (F) Spearman correlation between mRNA abundance (TPM) and the number of Ψ sites in a single mRNA, normalized by gene length. The bottom plot shows the Spearman correlation between mRNA abundance and Ψ levels (Δ deletion fraction). NS, not significant.

Next, we analyzed the microbiome composition and abundance using Kraken 2/Bracken^30^. We initially determined, using publicly available paired metagenomics and metatranscriptomics oral samples, that Kraken 2/Bracken accurately recapitulates microbial taxonomic information from metatranscriptome data (Extended Data Fig. 5A). In our patient samples, the microbiome was mostly dominated by Gram-negative bacteria, with low *Streptococcus* abundance, in agreement with the known microbial profiles of periodontitis patients^31,32^ (Fig. 6B). In total, we identified 174 Ψ sites from 141 transcripts (ranging from 1 – 4 sites per mRNA), distributed across 79 different genera and 11 phyla (Fig. 6C and D, Extended Data Fig. 5B, Supplementary Table 7). Strikingly, over 90% of pseudouridylated transcripts have unknown function (Fig. 6C, Supplementary Table 8). We also looked at the sequence preference for Ψ deposition and identified 106 different motifs (Fig. 6E, Supplementary Table 7). Although we were unable to assign most motifs to a PUS enzyme, the most abundant motif clearly resembles the TruB recognition sequence (Fig. 6E, Fig. 3E). Therefore, TruB is likely the major PUS enzyme in the oral microbiome.

Finally, we examined the relationship between Ψ and gene expression in the community. We found the degree of pseudouridylation (i.e. deletion fraction) did not correlate with mRNA level. Rather, the number of modification sites in an mRNA is positively correlated with transcript abundance (Fig. 6F). This phenomenon was observed in both patients’ samples (Fig. 6F). Therefore, multiple Ψ sites may be required to stabilize an mRNA.

Altogether, Ψ modifications are prevalent in microbial communities and our approach provided an avenue to study posttranscription regulation in microbiomes.

## Discussion

The lack of a suitable approach to study pseudouridylation in prokaryotes has made it difficult to identify exact modification sites in bacteria mRNAs. While a previous study could only identify modifications in 42 *E. coli* mRNAs^11^, our findings show that one-third of the *E. coli* transcriptome harbor Ψs. Moreover, our results provide a quantitative measure of Ψs. Therefore, our work represents a comprehensive analysis of bacterial mRNA pseudouridylation.

Although it is well established that Ψ stabilize rRNAs and tRNAs, there has been conflicting information regarding their role in mRNA stability. In one study, no association was found between Ψ levels and mRNA abundance in human HEK293T cells^15^ whereas, Dai *et al.* observed a significant association with mRNA levels^13^. In contrast, Nakamoto *et al* found modifications by TruA homolog (PUS1) resulted in transcript instability in *Toxoplasma gondii*^33^. Nevertheless, our data suggests Ψ deposition is significantly associated with mRNA levels in *E. coli*.

One major advantage of Ψ profiling is the ability to identify genes differentially expressed between two conditions which ordinarily may not be identified using traditional RNA-seq. This is particularly true for low-expression genes which can make it harder to detect differentially abundant transcripts^34^. In our data, mRNAs that are usually present in low abundance in *E. coli,* but slightly elevated in stress conditions, were solely detected using Ψ mapping. Therefore, the combination of the highly sensitive BS-based Ψ profiling and conventional RNA-seq can provide a more robust approach to study differential gene expression in prokaryotes.

Our data further indicates that mapping of Ψ positions can assist in the discovery of previously unidentified transcription units. In bacteria species where asRNAs have been studied so far, they can comprise almost 50% of the entire transcripts in a cell^35^. Yet, asRNAs which may potentially play a regulatory role in bacteria, has only been studied in a few microbes^35^. Methods to uncover novel transcripts, especially asRNAs, typically rely on sequencing-based approaches such as differential RNA-seq (dRNA-seq)^36^ coupled with strand-specific sequencing which preserves strand orientation^27^. However, our approach does not require strand information throughout the experimental pipeline.

Although previous studies in *E. coli* have identified transcription from the antisense strand of multiple tRNAs^27^, none have been identified so far from tRNA-Tyr. Transfer RNA-derived small RNAs have been shown to regulate gene expression in eukaryotes, and they are also implicated in the pathogenesis of many diseases^37^. It is plausible that sRNAs derived from reverse strands of tRNA may also play a general role in bacteria gene regulation. Interestingly, our candidate sRNA is found next to a characterized regulatory sRNA and a small open reading frame (ORF)^38,39^, suggesting this region may be a hotspot for regulatory genes.

In addition, there are currently no studies investigating posttranscription regulation in the microbiome. This is important because many pathogens use posttranscription regulation to control the expression of key virulent genes to adapt to their environment^40^. Likewise, we demonstrate that our approach can be extended to interrogate pseudouridylation in complex microbial communities. Similarly to our *E. coli* findings, we observed a correlation between Ψ sites and steady-state mRNA levels in the oral microbiota. As a result, the association between Ψ and transcript abundance may be a widespread phenomenon in prokaryotes.

In conclusion, our work provides a quantitative landscape of Ψ in *E. coli* and describes the interrogation of Ψ in microbiota samples. A major limitation of our approach is the high coverage requirement to call a Ψ position. This is evident in our inability to detect more Ψ sites in the microbiome samples. Similarly, when the Ψ site is next to multiple consecutive uridines, it is computationally challenging to determine the exact pseudouridylation site. Nonetheless, our findings have demonstrated the value of BS-based methods in the study of this important modification in bacteria. Future work on Ψ will not only expand our basic understanding of posttranscription regulation, but also provide new insights into the role of pseudouridylation on protein translation landscape.

## Methods

### Bacterial strains and growth conditions

Unless stated otherwise, wild-type (MC415), Ψ^ΔrRNA^ (MC452), Δ*truA*, Δ*truB*, Δ*truC*, and Δ*truD* mutants were routinely grown on Luria-Bertani (LB) media at 37°C (Supplementary Table 9). When required, 25 μg/ml of kanamycin was added to the media of Δ*tru* mutants. To induce stress conditions, bacteria (MC415 and MC452) were grown in the presence of ampicillin (0.4 μg/μl), gentamicin (0.4 μg/μl), or high salinity stress conditions (4% NaCl). In the latter, cells were cultured in Tryptic soy broth (TSB). In double-plasmid expression systems, cells were grown in M9 minimal media and gene expression from *lac* promoter was induced with 0.5 mM IPTG. Antibiotics were used at the following concentrations: 100 μg/ml carbenicillin; 50 μg/ml spectinomycin.

### RNA extraction, bisulfite treatment and library preparation

All fresh cultures were grown until OD_600_ of 0.3 and subsequently preserved in RNA protect (Qiagen, #76506). Next, RNA was extracted from the cultures using RNeasy mini kit (Qiagen, #74104) and residual DNA was removed by DNA-free^TM^ DNA Removal Kit (Thermo Fisher Scientific, AM1906). After RNA quantification and quality analysis by Qubit Flex and Bioanalyzer, RNA samples were diluted with nuclease-free water to give approximately 400-700 ng in 22 μl of volume which was thereafter split in two halves (i.e ‘BS-treated’ and ‘untreated’ halves). Ribosomal RNA was depleted in both halves using Ribo-Zero Plus rRNA Depletion Kit (Illumina, #20040525) followed by fragmentation of all samples by addition of 0.9 μl of fragmentation reagents (Thermo Fisher Scientific, #AM8740). The mix was incubated at 95°C for 20 s and fragmentation was stopped using 0.9 μl of stop reagent. Samples were immediately placed on ice.

Fresh bisulfite reagent (BSR) was prepared by adding 0.27 g of sodium sulfite (Sigma Aldrich, #901916) and 0.034 g of sodium bisulfite (Sigma Aldrich, 799394) to 900 μl of DEPC treated water. Next, 45 μl of freshly prepared BSR was added to the 11 μl of fragmented ‘BS-treated’ half and incubated at 70°C for 3 h. After incubation, 75 μl of nuclease-free water was added to the mix followed by 270 μl of RNA binding buffer (RNA Clean and Concentrator-5 column kit, Zymo Research, R1015) and 400 μl of 100% ethanol. The entire mixture (∼800 μl) was loaded onto the column and centrifuged for 30 s. The column was then washed with 200 μl of RNA wash buffer and subjected to desulfonation by adding 200 μl of RNA desulphonation buffer (Zymo Research #R5001-3-40) to the column. This was subsequently incubated at room temperature for 75 min. The column was washed using RNA wash buffer and eluted in 10 μl of elution buffer.

For the other ‘untreated’ half, samples were purified after fragmentation and eluted with 10 μl of elution buffer. Both purified BS-treated and untreated RNA samples were then annealed to random hexamers (50 μM) and subjected to cDNA first strand synthesis using SuperScript™ IV reverse transcriptase (Thermo Fisher Scientific, #18090010). The reaction parameters were: 23°C for 10 min, 50°C for 1 h, and 80°C for 10 min. Next, the second strand of cDNA was synthesized by DNA polymerase 1 (Thermo Fisher, #EP0041) in a reaction mixture containing 0.8 μl of RNAse H (Thermo Fisher, #EN0202) followed by incubation at 15°C for 2 hr and thereafter 75°C for 10 min. This double stranded cDNA was purified by adding 90 μl of Ampure beads (Beckman Coulter, A63881) using exact instructions provided by Illumina (Illumina Stranded Total RNA Prep, Ligation with Ribo-Zero Plus reference guide). Subsequent steps for 3’ ends adenylation, anchor ligation and indexing were performed using the Illumina Stranded Total RNA Prep, Ligation with Ribo-Zero (Illumina, #20040525), anchor plate (Illumina, #20040899), and DNA/RNA UD indexes set (Illumina, #20091646), respectively. Generated libraries were evaluated using Bioanalyzer and sequenced on the NovaSeq platform.

### Plasmid cloning and strain construction

To construct plasmids expressing sRNA, the pCDFDuet-1 vector was linearized via PCR using Q5® High-Fidelity DNA Polymerase (New England Biolabs, NEB) with primers listed in Supplementary Table 10. Fragments encoding sRNA with either the native promoter or the *lac* promoter was amplified from MG1655 genomic DNA via PCR. These fragments were separately inserted into the linearized pCDFDuet-1 vector using Gibson Assembly (NEB #E2611).

For constructing plasmids expressing superfolder GFP (sfGFP) under the control of candidate promoters, the promoter regions of the *sucA* and *rpoB* genes were amplified from MG1655 genomic DNA via PCR. The sfGFP coding sequence, synthesized by Twist Bioscience, was combined with each promoter and inserted into the linearized PHERD30Tamp vector using Gibson Assembly. An empty plasmid lacking a promoter was included as a control.

All plasmids were cloned into NEB® 5-alpha competent *E. coli* cells (#C2987H) and validated by whole-plasmid sequencing performed by Psomagen. Each sRNA-expressing plasmid was co-transformed with a corresponding promoter-GFP plasmid into *E. coli* MG1655 via electroporation.

Oligonucleotides used in this study were purchased from Integrated DNA Technologies and are detailed in Supplementary Table 10.

### Human oral sample collection

Collection of human samples was performed on an IRB approved clinical protocol at the NIH Clinical Center (ClinicalTrials.gov ID NCT01568697). All study participants provided written informed consent for participation in this study. Participants were deemed systemically healthy based on detailed medical history and select laboratory work up. In addition to systemic screening, periodontal status was assessed through detailed clinical oral evaluation. Subgingival plaque samples (tooth adherent biofilm) were removed using a Gracey Curette (HuFriedy Group), from two patients with severe periodontitis (AAP, stage IV). Subgingival plaque samples were immediately placed in DNA/RNA Shield Stabilization Solution (Zymo Research, #R1100-50).

### RNA extraction from human samples

Samples were vortexed to dissolve pellets and then added to 2ml ZR BashingBead Lysis Tubes (0.1 & 0.5 mm) (Zymo Research, #S6012-50) with 600 μl of Qiazol lysis solution (Qiagen, #79306). Bead beating was performed for 5 min in FastPrep 24 homogenizer at maximum speed. After centrifugation, 180 μl of chloroform was added to the supernatant and centrifuged for another 15 min at 4°C for phase separation. The aqueous phase was transferred to a new tube and 1.5:1 v/v 100% ethanol was added. The mixture was mixed by pipetting and RNA was extracted and processed as described in the previous section.

### Bioinformatics pipeline to detect Ψ

Adapter removal and quality trimming of raw reads was performed using Trim Galore^41^. We mapped clean reads to *E. coli* BW25113 genome (NZ_CP009273.1) using bwa-mem^42^ and realigned reads using ABRA2 to improve detection of indels in downstream analysis^16^. Next, bam-readcount was used to retrieve nucleotide coverage and sequence variant information^43^ and the resulting output was subsequently parsed using brc-parser.py^44^. To be considered a Ψ site, a uridine position needs to meet the following criteria: (i) coverage depth ≥ 20 in both BS-treated and untreated samples; (ii) deletion count ≥ 5 in BS-treated samples; and (iii) deletion fraction ≥ 5% in BS-treated samples but less than 1% in untreated samples. A Ψ site was considered as high confidence if it was identified in at least 2 samples in the main dataset (Extended Data Fig. 1A).

### Motif analysis

To determine the sequence preference for each PUS, we retrieved flanking sequences of mRNA sites having deletion fraction > 6%, which is the median deletion fraction in our dataset. We further excluded low occurrence motifs which were identified less than 5 times. For every motif passing the threshold, deletion fractions were compared between WT and Ψ^ΔrRNA^ strains, or between the four different *tru* mutants. A motif significantly decreased (*P* value < 0.05, two-sided Wilcoxon rank sum test) in a mutant was considered a sequence preference for that PUS. To identify consensus sequences, motifs were submitted to MEME and the parameter to search the reverse complement strand was disabled^45^.

### Pathway analysis

The pathways of genes associated with pseudouridylation were analyzed using the KEGG Genes database for *E. coli*. The KEGG Mapper search tool was employed to map and assign these genes to the corresponding pathways^46^.

### RNA structure analysis

tRNA structures were predicted with RNAfold web server using default parameters^47^. To investigate base-pairing preference for pseudouridylation, 12-mer sequences flanking each side of the Ψ residue were used as input in the command-line version of RNAfold (-p -T 37 parameters).

### Estimation of mRNA abundance from RNA-seq and influence of Ψ on stability

Because untreated samples are equivalent to traditional RNA-seq, we quantified the mRNA abundance from read counts in untreated samples. To begin, the generated .bam files from previous mapping step were used as input in featureCounts^48^ to retrieve unique read counts mapped to each gene. We included a pseudocount of one to each gene and normalized using ‘transcript per million’ (TPM).

To evaluate the role of Ψ on mRNA abundance, we initially calculated the Ψ-strength for each transcript as previously described^13^. Ψ-strength is defined as the sum of deletion fractions at all Ψ sites within one RNA. The gene-level deletion fractions were compared between WT and PUS mutants using only samples obtained at 37°C because *tru* mutants were solely grown at optimum temperature. For a gene to be considered pseudouridylated by a given PUS, the difference in gene-level Ψ-strength between WT and a mutant must be ≥ 0.05 in both biological replicates.

### Determination of differential pseudouridylation in stress conditions

To identify transcripts with increased Ψ under stress conditions, we calculated the Ψ-strength for each mRNA. Similarly, an mRNA was considered to harbor increased Ψ if the difference in gene-level Ψ-strength between a stress condition and control (37°C) samples was ≥ 0.05 in both biological replicates. We further performed manual inspection to exclude false positives potentially arising from candidates that narrowly missed the cutoff in control samples.

### RT-qPCR

Primers were designed using PrimerQuest^®^ to give ∼ 100 bp amplicons (Supplementary Table 10). Extracted RNA was first normalized to ensure similar RNA concentration in each reaction.

Synthesis of cDNA was performed using high-capacity cDNA RT kit (Thermo Fisher, #4368814) prior to qPCR reaction using SSoAdvanced Universal SYBR Green Supermix (Biorad, #1725270).

For every group, four biological replicates were used. For each sample, technical duplicates were performed and the *Ct* values were normalized to 16S rRNA gene (Δ*Ct*). Statistical significance was determined from normalized *Ct* values using unpaired, two-tailed *t*-test and relative fold expression was calculated using 2^−Δ*Ct*^ method^49^.

### Antisense-RNA analysis

The opposite strand for each gene was examined for the presence of Ψ. The predicted asRNA were compared to previously reported asRNAs in Thomason *et al*^27^. To predict promoter sequences upstream (200 bp) of Ψ sites, we used the software iProEP^28^ and only considered outputs with a probability score ≥ 0.9. The subsequence with the highest probability was considered as the region containing the promoter. The tool SAPPHIRE^50^ was subsequently used to estimate the TSS.

For antisense small RNA analysis, we predicted transcription termination sites from 150 bp sequences downstream the Ψ site using iTerm-PseKNC^51^. Similarly, results were filtered using a probability cutoff of ≥ 0.9. Target sites of putative sRNAs were predicted with TargetRNA3 (0.75 probability)^52^ using sequences starting from TSS (Supplementary Table 5). Length of putative sRNAs were predicted to span from TSS to midpoint in subsequence containing the TTS.

### Microbiome analysis

We downloaded 3,589 oral metagenome-assembled genomes (MAGs) and their corresponding taxonomic information from Zhu *et al.*^29^, predicted genes using Prodigal (-p meta)^53^, and clustered sequences at 90% identity using MMseqs2 (--min-seq-id 0.9 -c 0.9 --cov-mode 1 --cluster-mode 2)^54^. Reads were mapped to these clusters using bwa-mem^42^, realigned using ABRA2^16^, and per-base coverage was retrieved using bam-readcount^43^. Uridine positions passing the following filters were considered as pseudouridylated: (i) read depth ≥ 20 in both BS-treated and untreated samples; (ii) deletion count ≥ 5 in BS-treated samples; (iii) deletion fraction ≥ 0.02 (i.e. 2%) in BS-treated samples; (iv) deletion fraction in BS-treated samples is, at least, 2 fold above deletion fraction in untreated samples; and (v) the *P*-value from Fisher’s exact test should be < 0.01 when total number of deletions and reads depth in BS-treated and untreated samples are compared. For sites passing the above threshold, the proportion of Ψ was derived as the difference in deletion fraction between BS-treated and untreated sites (i.e. Δ deletion fraction).

Transcript abundance was estimated similarly to *E. coli* isolate data. However, we excluded genes with less than 5 reads in all samples prior to TPM calculation. Functional annotation of transcripts was performed using eggNOG^55^. Finally, to evaluate the accuracy of Kraken 2/Bracken in metatranscriptomics data, we retrieved paired metagenomics and metatranscriptomics oral samples from Belstrøm *et al*^56^.

## Data availability

Raw sequence reads generated in this study will be deposited in Sequence Reads Archive (SRA) upon acceptance. For paired oral metagenomics and metatranscriptomics analysis (BioProject PRJNA396840), the following SRA accessions were used; SRR5892221, SRR5892220, SRR5892219, SRR5892218, SRR5892225, SRR5892224, SRR5892223, SRR5892222, SRR5892227, SRR5892226, SRR5892199, SRR5892198, SRR5892197, SRR5892196, SRR5892203, SRR5892202, SRR5892201, SRR5892206, SRR5892194, SRR5892193. All other necessary data are included in the Supplementary Information.

## Acknowledgements

This work was funded by the Intramural programs of the NIDCR and NIAID. We are grateful to NIDCR/NIDCD Genomics and Computational Biology Core (ZIC DC000086) for providing sequencing support and Dr. Michael O’Connor for sharing *E. coli* strains MC415 (WT) and MC452 (Ψ^ΔrRNA^). We also thank Laurie Brenchley and Teresa Wild for obtaining and processing subgingival plaque samples.

## Author contributions

S.S and A.E conceived the study. S.S, B.W, B.Y, and M.P performed wet-lab experiments. A.E and N.D. conducted bioinformatics analyses. S.S, A.E, and N.M analyzed the data. S.S. and A.E. wrote the manuscript. A.E supervised the study.

---

**Extended Data Figure 1:**
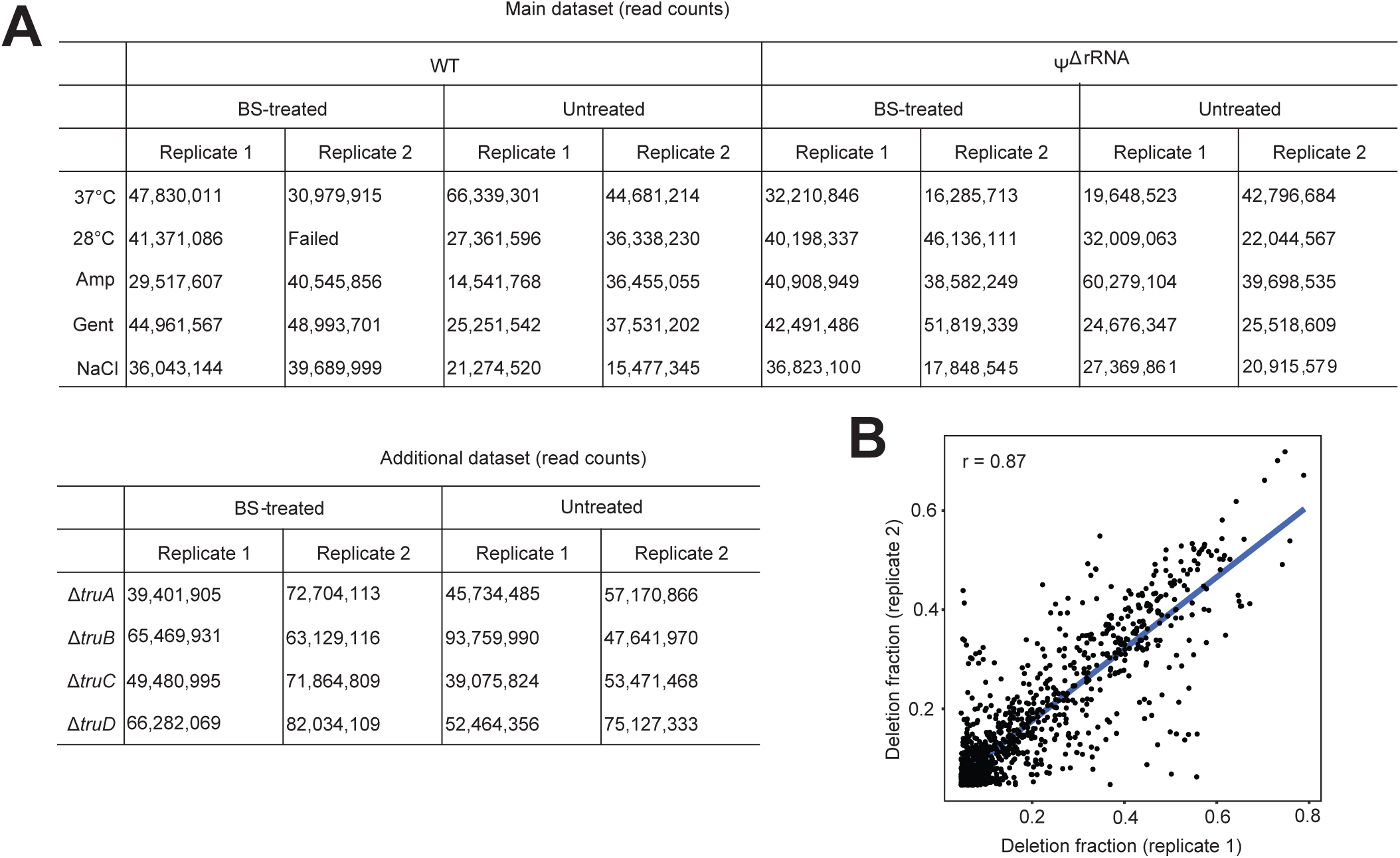
Sequencing data and quantitative reproducibility of BS-based Ψ profiling. (A) Total read counts of WT and mutant strains. (B) Pearson correlation between deletion fractions from two biological replicates.

**Extended Data Figure 2:**
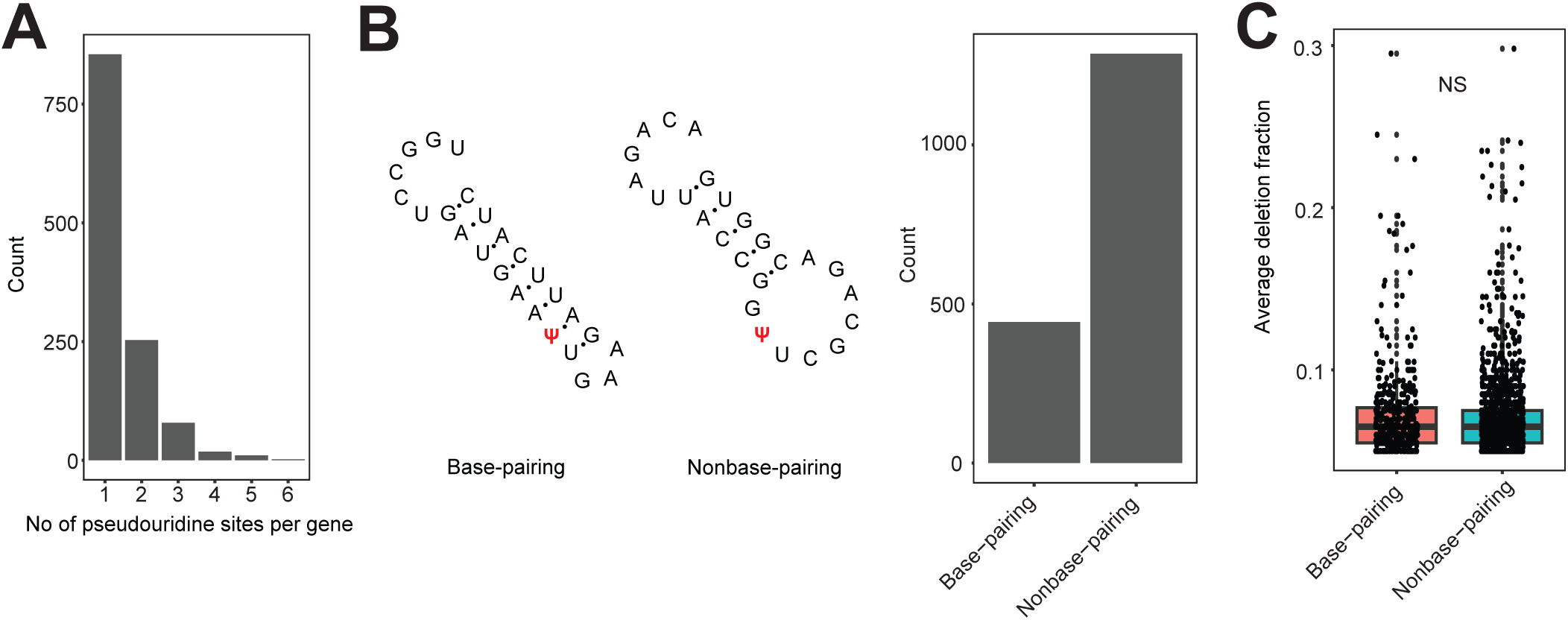
Role of mRNA secondary structure on pseudouridylation. (A) Distribution of the number of Ψ sites found in a single mRNA. (B) Examples of base-pairing and nonbase-pairing Ψ sites. The Barplot shows the distribution of pseudouridylation in paired or unpaired uridine residues. (C) Boxplot showing average deletion fractions in paired and unpaired Ψ sites. NS, not significant.

**Extended Data Figure 3:**
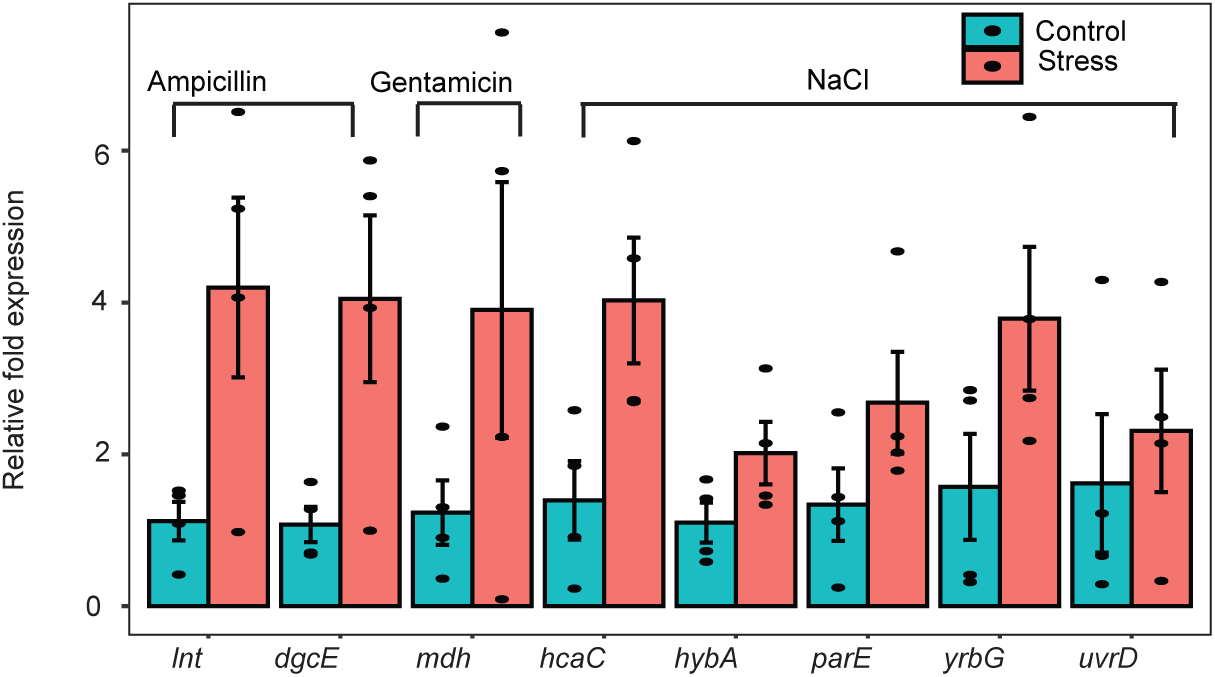
mRNA quantification using RT-qPCR. Relative levels of mRNAs that are not enriched in stress conditions based on statistical significance (n = 4). Error bars represent SEM.

**Extended Data Figure 4:**
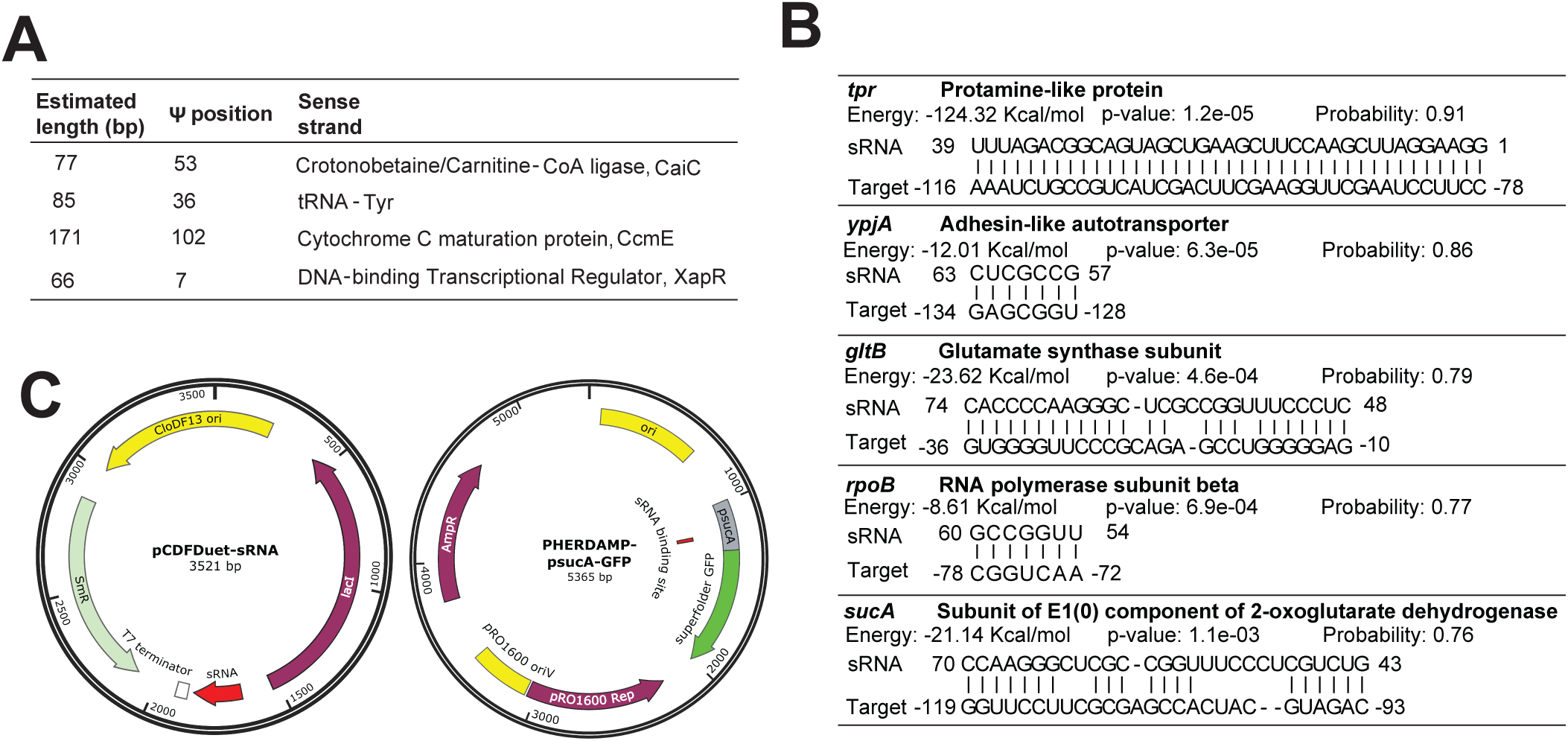
Characterization of small RNAs identified by Ψ mapping. (A) Information on putative sRNAs transcribed from antisense strands. (B) Target sequences for sRNA transcribed from the antisense strand of tRNA-Tyr. Predictions were made with TargetRNA3. Because *tpr* is located close to tRNA-Tyr, the upstream sequence shown is not the UTR, but rather the sequence of tRNA-Tyr. (C) Plasmid maps used to construct the double-plasmid expression system.

**Extended Data Figure 5:**
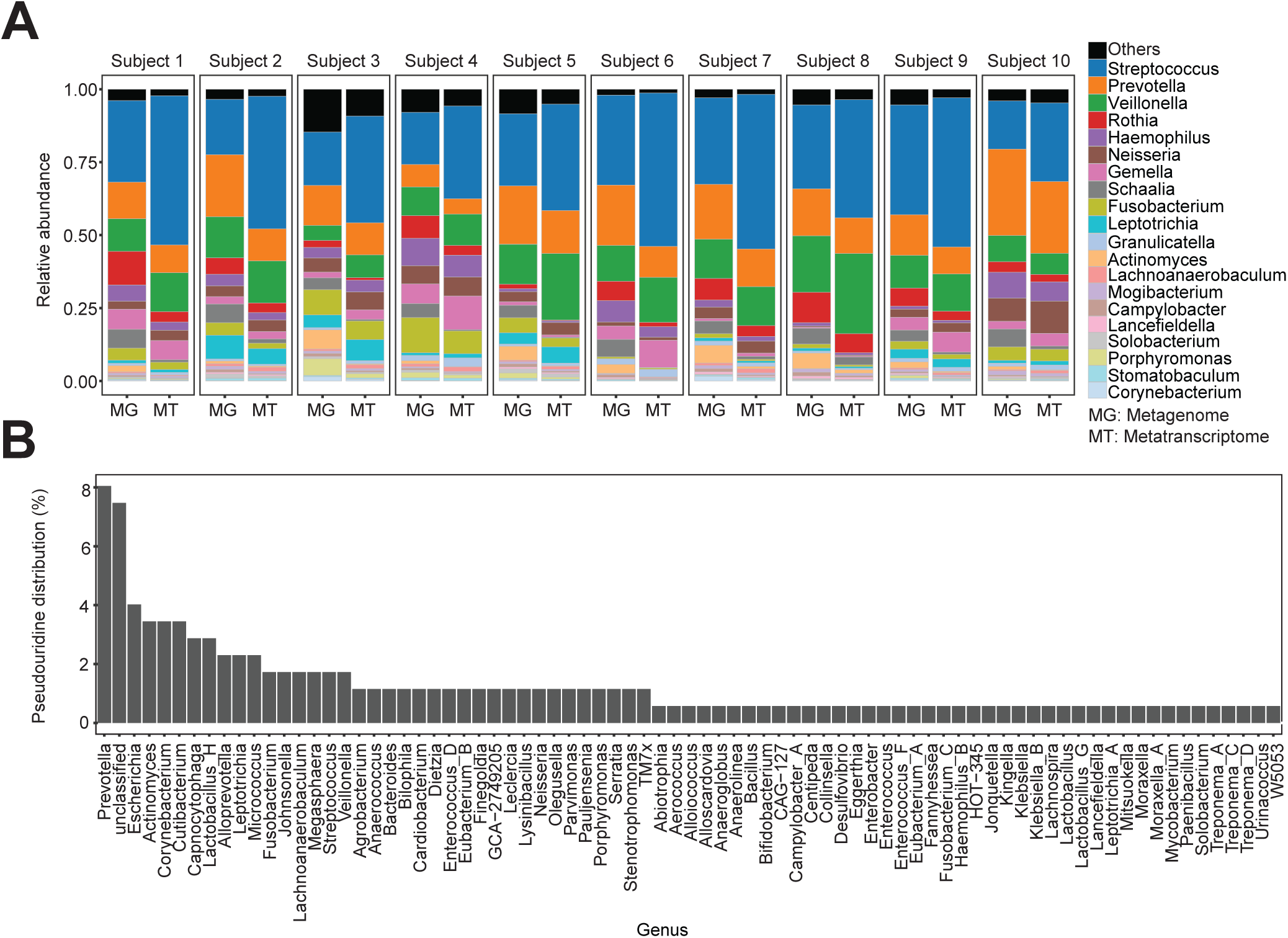
Oral microbiome analysis. (A) Comparison of microbial profiles in paired metagenomic and metatranscriptomics oral samples (Belstrøm *et al.*) using Kraken 2/Bracken. (B) Distribution of pseudouridylated mRNAs across all bacteria genera.

## Supplementary Tables

**Table S1: Pseudouridine profiles in WT and ΨΔrRNA *E. coli* strains under different growth conditions**

**Table S2: Pseudouridine profiles in *tru E. coli* mutants**

**Table S3: Sequence motifs for pseudouridylation**

**Table S4: Pseudouridine profiles from antisense genes**

**Table S5: Predicted small RNAs from antisense genes**

**Table S6: Nucleotide sequences for sRNA experimental validation**

**Table S7: Pseudouridine profiles in oral microbiome samples**

**Table S8: Oral microbial genes with annotations**

**Table S9: Plasmids and bacteria strains used in this study**

**Table S10: Primers used in this study**

